# Genome of *Trichoderma gamsii* strain T035 a promising beneficial fungus in agriculture

**DOI:** 10.1101/2025.09.22.677720

**Authors:** Justine Colou, Etienne Brémand, Alwena Robin, Franck Bastide, Thomas Guillemette

## Abstract

*Trichoderma gamsii* is a filamentous fungus widely recognized for its beneficial roles in agriculture, particularly for its ability to suppress plant pathogens and enhance crop health. However, genomic resources for this species remain scarce, limiting functional and applied studies. Here, we report the high-quality genome of *T. gamsii* strain T035, a promising biocontrol strain with significant antagonistic activity against several pathogens *in vitro*. The assembly consisted of 16 sequences, including near 7 chromosome-scale sequences, with an N50 value of 7.2 Mbp and a total assembly length of 38.8 Mbp. This genome represents the most complete *T. gamsii* assembly to date and will provide a valuable resource to facilitate the exploration of molecular mechanisms underlying biocontrol and support the development of sustainable plant protection strategies.

## Introduction

The genus *Trichoderma* (Ascomycota, Hypocreaceae) includes fungi widely recognized for their biocontrol potential in agriculture. To date, 225 *Trichoderma* genomes are publicly available on NCBI, but only a few have been assembled to completion: the reference genomes of *Trichoderma atroviride, Trichoderma asperellum, Trichoderma asperelloides, Trichoderma simmonsii*, and *Trichoderma semiorbis*. Complete genomes are also available for *Trichoderma reesei* and *Trichoderma virens*, although these are not designated reference genomes. High-quality assemblies are particularly valuable for comparative genomics and for identifying genes involved in pathogen suppression, secondary metabolism, and other biocontrol-related functions (Li *et al*., 2021; Rosolen *et al*., 2022; Schalamun et Schmoll, 2022).

*Trichoderma gamsii* is a ubiquitous fungus found in diverse environments, where it can exhibit psychrotolerance and broad pH tolerance (Migheli *et al*., 2009; Rinu, Sati et Pandey, 2014). This species is particularly interesting due to its ability to suppress pathogens through enzymatic activity and the release of volatile organic compounds (Anees *et al*., 2010; Chen *et al*., 2016; Bedine Boat *et al*., 2020; Galletti, Paris et Cianchetta, 2020; Valan Arasu *et al*., 2023; Marik *et al*., 2025). Additionally, it promotes plant health by inducing systemic resistance and stimulating plant growth (Anees *et al*., 2010; Rinu, Sati et Pandey, 2014; Chen *et al*., 2016; Galletti, Paris et Cianchetta, 2020; Valan Arasu *et al*., 2023). These characteristics highlight the potential of *T. gamsii* for biotechnological applications, underscoring the importance of comprehensive genomic data for further investigation of its biocontrol abilities. To date, the best public genomic sequence available for this species belongs to the strain T6085, which is known for its ability to control *Fusarium* head blight on wheat (Baroncelli *et al*., 2016).

In this study, we performed *de novo* sequencing of *T. gamsii* T035 genome. This strain exhibits strong antagonistic activity against multiple plant pathogens, making it a promising model to explore *T. gamsii* biocontrol potential. Here, we present a high-quality, near chromosome-level genome assembly and its functional annotation. With only 16 sequences, seven of which corresponding to near chromosome-scale nuclear scaffolds, this assembly represents the most contiguous *T. gamsii* genome available to date and provides a valuable resource to support future comparative and applied studies.

## Material and methods

### Sample information

*Trichoderma gamsii* strain T035 was isolated in 2019 from carrot-cultivated soil in Brain-surl’Authion, France (GPS coordinates: latitude 47.473936; longitude −0.390684) (Fig.1). Sampling and isolation were performed following the method described by Chateau et al. (2024). Briefly, 5 g of rhizosphere soil were suspended in 50 mL of sterile water. Serial dilution–extinction was carried out on Potato Dextrose Agar (PDA; Grosseron, 39 g L^−1^) supplemented with streptomycin (500 mg L^−1^). Cultures were incubated in darkness at 20 °C for seven days. Strain T035 was subsequently transferred to Malt Agar medium (MA; Grosseron, 20 g L^−1^ bacteriological malt extract, 15 g L^−1^ bacteriological agar type A) and maintained under the same growth conditions. For long-term preservation, mycelial explants were stored at −80 °C in 30% glycerol solution as a cryoprotectant.

**Figure 1.**
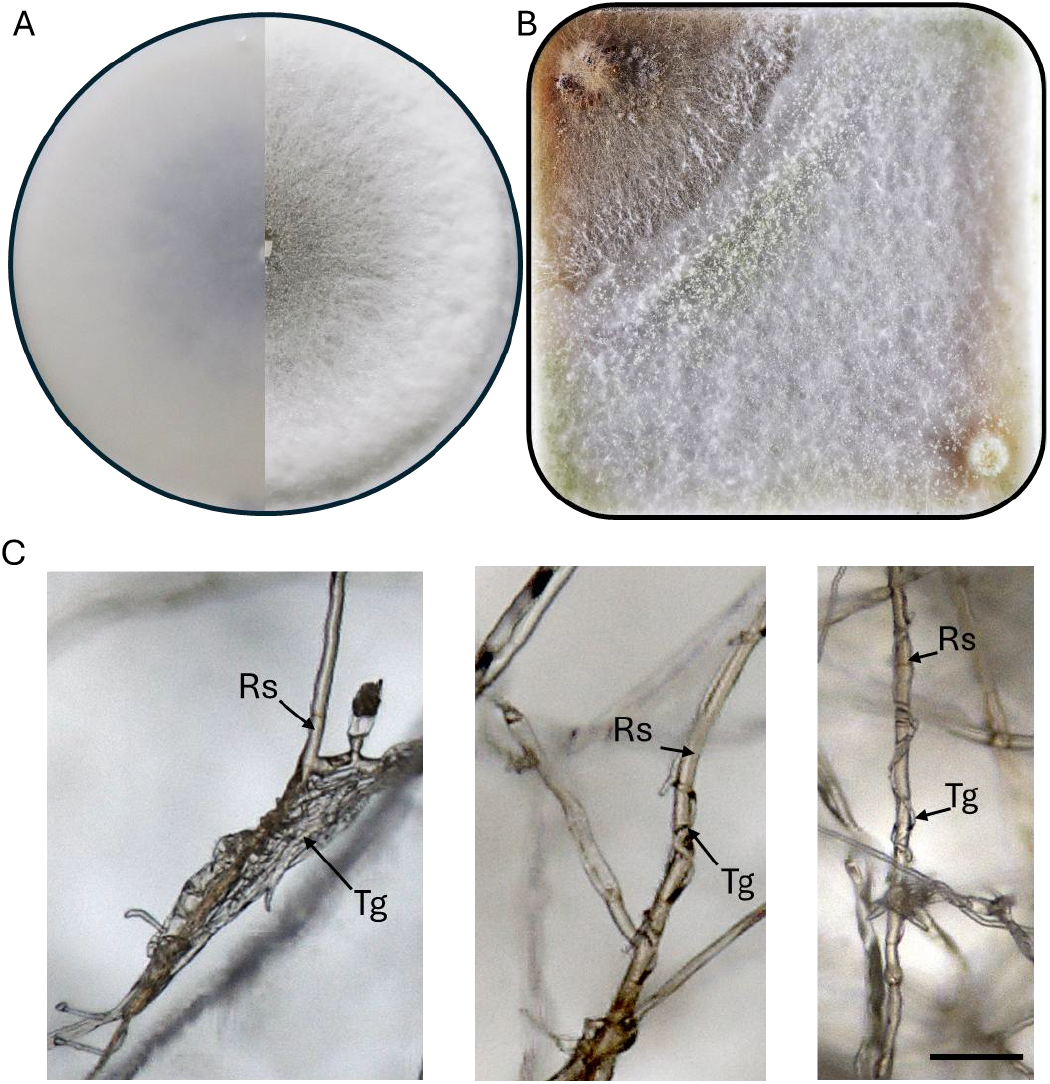
Colony morphology of *Trichoderma gamsii* T035 on PDA after 7 days of incubation, with bottom (left) and top (right) views of the plate (A). Confrontation assay between *T. gamsii* T035 (bottom-right) and *Rhizoctonia solani* PAT-009 (top-left) (B). Hyphal coiling of *T. gamsii* T035 (Tg) around *R. solani* PAT-009 (Rs) (scale bar = 50 µm) (C).

*Trichoderma atroviride* strain I1237 was obtained from the commercial product TRI-SOIL® (Agrauxine, Beaucouzé, France).

Pathogenic strains were sourced either from our laboratory fungal collection (IRHS, FungiSem Team, Beaucouzé, France) or from the CIRM-CFBP collection (Beaucouzé, France). The twelve filamentous pathogens belong to *Rhizoctonia solani* (PAT-002/PAT-006/PAT-009), *Globisporangium* sp. (PAT-013/PAT-018/PAT-019), *Stagonosporopsis valerianellae* (PAT-023/PAT-024/PAT-025), and grapevine trunk disease fungi, namely *Phaeomoniella chlamydospora* (PAT-026), *Diplodia seriata* (PAT-027), and *Neofusicoccum parvum* (PAT-028). The two bacterial strains used in this study were *Acidovorax valerianellae* (PAT-034) and *Xanthomonas campestris* (PAT-035).They are further described in the Supplementary Table S1.

### *In vitro* control of filamentous pathogens

The two *Trichoderma* isolates were evaluated for their mycoparasitic activity against the twelve pathogenic fungal and fungal-like isolates previously described using the dual culture assay (Bunbury-Blanchette et Walker, 2019). Square Petri dishes (120 × 120 mm) containing PDA medium were inoculated with two 5 mm mycelial plugs: one obtained from a pathogen colony and the other from a *Trichoderma* colony previously grown on PDA under the conditions described above. The plugs were placed at opposite corners of the plate, with the pathogen inoculated two days before the antagonist. Each condition was replicated three times, and the entire experiment was repeated independently three times.

Plates were incubated in darkness at 20 °C for 20 days. Mycoparasitic ability was visually assessed using a scale adapted from Pascouau et al. (2023): 0 = <5% of pathogen colony overgrown by *Trichoderma*; 1 = 5–25% overgrown; 2 = 25–50% overgrown; 3 = 50–75% overgrown; 4 = >75% overgrown (Fig. S1).

Statistical analyses were performed in R v4.3.1. Data were analyzed using a non-parametric analysis of variance (Wilcoxon signed-rank test), and *p*-values were adjusted for multiple testing using the false discovery rate (FDR) method.

### *In vitro* control of bacterial pathogens

Two *Trichoderma* isolates were assessed for their antibiosis capacity against the bacterial isolate PAT-034 (*A. valerianellae*) and PAT-035 (*X. campestris*) following the protocols of Dennis & Webster (1971) and Lee et al. (2012), with modifications. For each *Trichoderma* isolate, 250 mL Erlenmeyer flasks containing 50 mL of Potato Dextrose Broth (PDB, Grosseron, 24 g L^−1^, pH 5.2) were inoculated with five mycelial plugs (5 mm diameter) taken from 7-day-old PDA cultures. Cultures were incubated at 20 °C on a rotary shaker (100 rpm) for 7 days and subsequently filtered through sterile 0.2 μm syringe filters (ClearLine®, Dutscher) to obtain culture filtrates.

The effect of the filtrates on bacterial growth was evaluated using a spectrophotometric microplate assay (96-well plates, SPECTROstar, BMG Labtech). Autoclaved Lysogeny Broth (LB, Grosseron, 24 g L^−1^) was inoculated with 1% (v/v) of a bacterial suspension adjusted to OD = 0.05, supplemented with 10% (v/v) of *Trichoderma* culture filtrates, in a final volume of 300 μL per well. Filtrates obtained from non-inoculated PDA disks were incubated under the same conditions and served as controls. Plates were incubated at 25 °C for 33 h with intermittent shaking (5 min at 200 rpm every 10 min), and absorbance at 600 nm was recorded every 10 min.

Each condition was tested in triplicate, and three independent experiments were conducted. The area under the growth curve was calculated as described by Joubert et al. (2010) and Warringer & Blomberg (2003). Data were analyzed using analysis of variance (ANOVA) followed by Tukey’s multiple comparisons test (α = 0.05).

### Genome-wide relatedness among Trichoderma species

A global comparison of *Trichoderma* genomes was performed using MashTree v1.2 (Katz *et al*., 2019). Genome assemblies of *T. gamsii* and representative assemblies from other *Trichoderma* species were retrieved from NCBI RefSeq (accessions listed in Supplementary Table S2).

Pairwise genomic distances were computed with Mash (genome size parameter set to 40 Mbp, sketch size = 20,000). The distance matrix was used to infer a neighbor-joining tree with MashTree. The obtained tree was exported in Newick format, and later visualized and annotated using MEGA12 (Kumar *et al*., 2024) (Fig.2).

**Figure 2.**
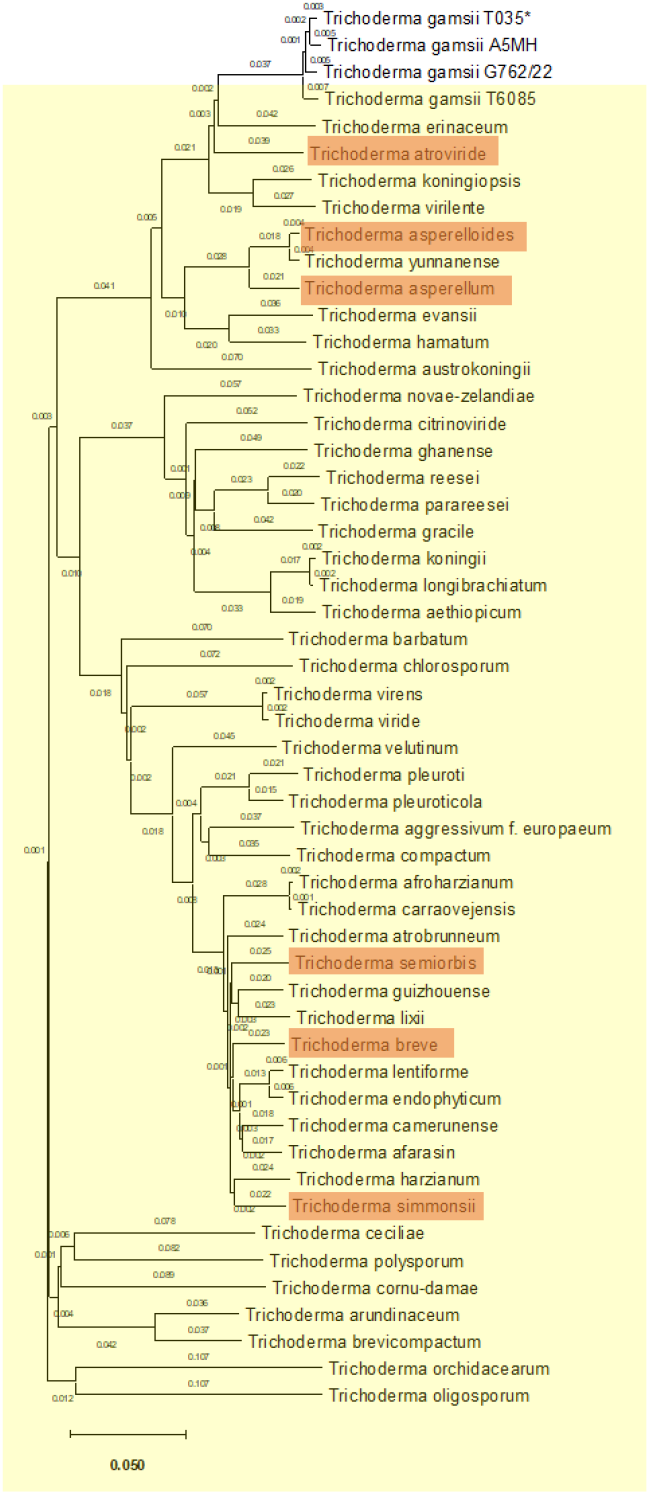
Genetic distance-based tree of *Trichoderma* genomes. The analysis included all reference *Trichoderma* genomes available in NCBI (yellow) as well as all known *T. gamsii* genomes. The genome obtained in this study is indicated with an asterisk. Complete genomes are highlighted in orange.

### DNA extraction

High-molecular-weight genomic DNA was extracted from a 7-day-old culture of *T. gamsii* T035 grown on a pectocellulosic membrane placed on PDA medium. Tissues were frozen with liquid nitrogen and bead-milled. DNA extraction followed an adapted Möller protocol: RNase treatment was performed immediately after proteinase K digestion, and DNA was resuspended in 10 mM Tris (pH 8) instead of TE buffer, as EDTA can interfere with Nanopore sequencing (Möller *et al*., 1992). The A260/280 and A260/230 ratios were 1.85 and 2.3, respectively. DNA concentration was measured using the quBit to 27 ng/µL, and its integrity was confirmed by electrophoresis gel, where the majority of the gDNA was above 10 kilobases (kbp) in length.

### Oxford Nanopore Technologies (ONT) sequencing

The sequencing library was prepared using the Native Barcoding Kit 96 V14 (SQK-NBD114.96) with 1.5 µg of high-molecular-weight genomic DNA as described in the kit protocol. We performed *de novo* sequencing of *T. gamsii* T035 genome using Oxford Nanopore Technologies’ MinION device M1kD on the flowcell FLO-MIN114 with R10.4 pores (Lu, Giordano et Ning, 2016).

### Genome assembly

Reads were basecalled using the super accuracy (SUP) mode. The long-read assemblies were performed using both Canu v2.2 (Koren *et al*., 2017) and Flye v2.9.6 (Kolmogorov *et al*., 2019). The less fragmented assembly was obtained using Flye, with reads filtered before assembly where reads shorter than 1,000 bp or with a quality score below Q10 were removed (De Coster *et al*., 2018). The assembly underwent three additional polishing steps using Racon v1.5.0 (Vaser *et al*., 2017). Contigs were scaffolded with RagTag v2.1.0 (Alonge *et al*., 2022) using *T. atroviride* GCF_020647795.1 as reference. Assembly quality was later assessed with BUSCO 5.4.4 (fungi_odb10 database) and compared to the reference *T. gamsii* genome (RefSeq: GCF_001481775.2) using QUAST v5.3.0 (Mikheenko *et al*., 2023; Tegenfeldt *et al*., 2025) (Fig. 3 and Table 1). Additionally, k-mer spectra plots were generated from Canu-corrected reads (Koren *et al*., 2017) using Jellyfish v2.2.10 (Marçais et Kingsford, 2011), and Genoscope 2.0 (Ranallo-Benavidez, Jaron et Schatz, 2020) (Fig. S2 - S5).

**Table 1.** Comparison of global genome metrics of *Trichoderma gamsii* genomes. The asterisk indicates the actual reference genome of *T. gamsii* in NCBI. The “m” indicates the mitochondrial sequences.

| ASSEMBLY STATISTICS | TRICHODERMA GAMSII STRAIN T6085* | TRICHODERMA GAMSII STRAIN A5MH | TRICHODERMA GAMSII STRAIN G762/22 | TRICHODERMA GAMSII STRAIN T035 (THIS STUDY) |
| --- | --- | --- | --- | --- |
| SUBMITTED GENBANK ASSEMBLY | GCA_001481775.2 | GCA_002894205.1 | GCA_049996675.1 |  |
| GENOME SIZE | 37.9 Mbp | 38.5 Mbp | 38 Mbp | 38.8 Mbp |
| TOTAL UNGAPPED LENGTH | 37.9 Mbp | 38.5 Mbp | 38 Mbp | 38.8 Mbp |
| NUMBER OF SCAFFOLDS | 172 | 292 | 499 | 8 (7+1<sup>m</sup>) |
| SCAFFOLD N50 | 697.4 kbp | 594.5 kbp | 285.3 kbp | 7.2 Mbp |
| SCAFFOLD L50 | 18 | 20 | 44 | 3 |
| NUMBER OF CONTIGS | 216 | 517 | 499 | 30 |
| CONTIG N50 | 534 kbp | 299.2 kbp | 285.3 kbp | 3.096 Mbp |
| CONTIG L50 | 23 | 42 | 44 | 5 |
| GC PERCENT | 49 | 48.5 | 48.5 | 48.4 |
| GENOME COVERAGE | 50x | 90x | 35x | 25.4x |
| ASSEMBLY LEVEL | Scaffold | Scaffold | Contig | Scaffold |

**Figure 3.**
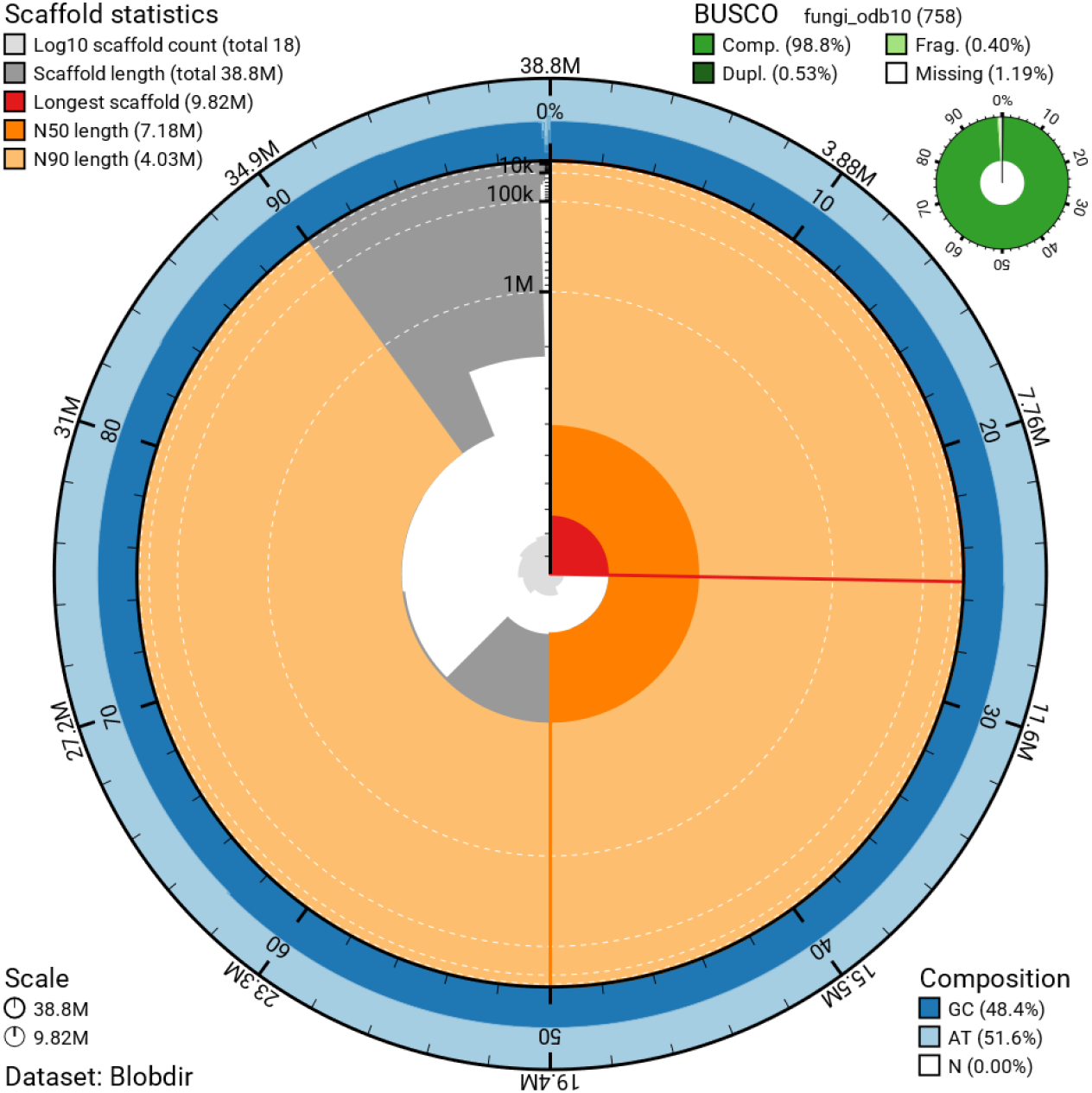
Snailplot generated with BlobToolKit showing assembly statistics for the *Trichoderma gamsii* T035 genome. The assembly has a total length of 38,792,561 bp with a maximum scaffold length of 9,818,134 bp (in red). N50 and N90 sequence lengths are 7,178,191 bp and 4,032,368 bp, indicated by orange and pale-orange arcs, respectively. The grey spiral represents scaffold lengths in descending order on a log scale. BUSCO analysis indicates 98.8% completeness (top right). The outer ring shows base composition (GC: 48.4%, AT: 51.6%, N: 0.0%). It’s important to note that BlobToolKit reports 0.0% N, as the gaps introduced by RagTag during the scaffolding are synthetic and not associated with a raw sequence.

### Annotation

Structural annotation was performed using the deep-learning tool Helixer (Holst *et al*., 2023). Telomeres were detected with TIDK v0.2.65, allowing the identification of 1,097 repeats of the canonical motif AAAAAAT (Hofstatter *et al*., 2022). To confirm telomere locations and identify centromeres, a sliding window analysis was performed using bedtools v2.31.1 and RepeatMasker v4.2.2 to detect regions with high repeat density, low GC content, and low gene density. Centromeric and telomeric regions were then manually curated by integrating overlapping signals from these analyses. The distribution of transposable elements (TEs) across the genome was assessed using EDTA v2.2.2 (Su *et al*., 2021) (Supplementary Table S3).

Functional annotation was carried out with EggNOG-mapper v2 (eggNOG database version 5.0.2) (Cantalapiedra *et al*., 2021) (Supplementary Table S4), signal peptides were predicted using DeepSig (Version 1.2.5) (Savojardo *et al*., 2018) (Supplementary Tables S5-6), secondary metabolite (SM) clusters were identified using antiSMASH 8.0 (fungal mode) (Blin *et al*., 2025) (Supplementary Table S7), and carbohydrate-active enzymes (CAZymes) were identified using dbCAN3 (dbCAN HMMdb v12) (Zheng *et al*., 2023) (Supplementary Table S8). Only high-confidence enzymes were kept in this study, defined as matches detected by all tested tools: HMMER: dbCAN (E-value < 1e-15, coverage > 0.35), DIAMOND: CAZy (E-value < 1e-102), and HMMER: dbCAN-sub (E-value < 1e-15, coverage > 0.35). All annotation results are synthesized in the Supplementary Table S9. A comparison of secondary metabolite clusters, proteins with signal peptide, and carbohydrate-active enzymes was performed on *T. gamsii* genomes as well as the complete reference genomes of *Trichoderma* available on NCBI (Supplementary Table S2).

### Trichoderma genome comparison

To identify syntenic regions between *T. gamsii* and *Trichoderma atroviride* (Fig. 6B), as well as among the different *T. gamsii* genomes (Fig. S6), comparative genomic and synteny analyses were performed using the JCVI toolkit v1.4.16 (Tang *et al*., 2024). Orthologous genes were identified based on protein sequence similarity using BLAST, and conserved syntenic regions were inferred from the genomic distribution and colinearity of these orthologs along chromosomes. Syntenic blocks were filtered to retain only robust regions supported by at least 30 consecutive colinear orthologous genes. Filtered syntenic relationships were visualized using JCVI-based karyotype representations.

## Results

### Biocontrol assays

The biocontrol potential of *T. gamsii* strain T035 was assessed through its parasitism and antibiosis capacities. Strain T035 displayed greater parasitic activity than the commercial strain I1237 against eight of the twelve tested filamentous plant pathogens (Fig. 4A). Notably, while I1237 exhibited antibiosis activity against both *Acidovorax valerianellae* and *Xanthomonas campestris*, strain T035 was more than twice as effective (Fig. 4B).

**Figure 4.**
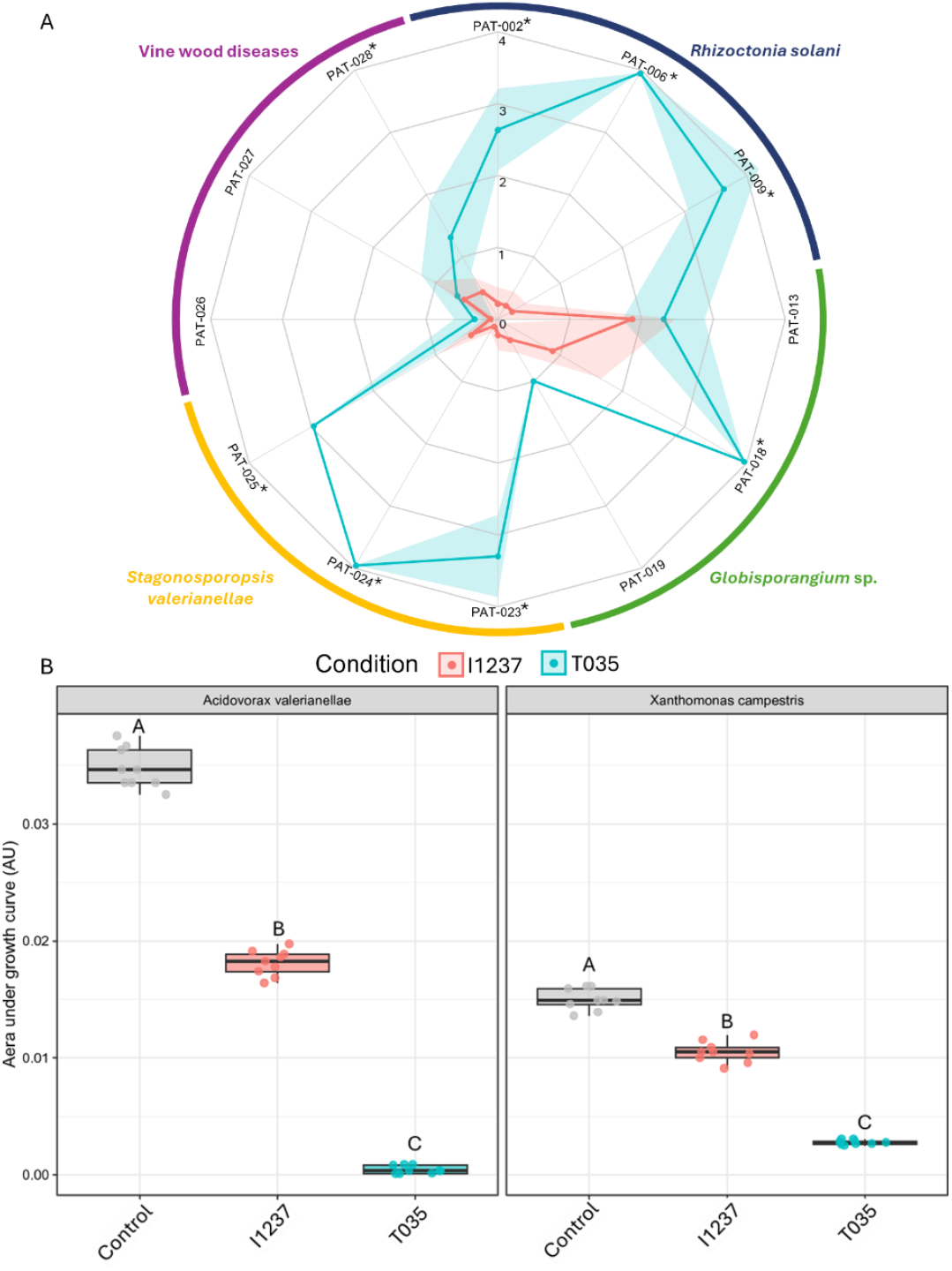
Evaluation of biocontrol capacities of *T. gamsii* strain T035 compared with a commercial *Trichoderma* strain (I1237). (A) Radar plot showing the parasitic capacity of both strains against twelve isolates from four different filamentous plant pathogens. Scores range from 0 (no colonization) to 4 (complete colonization). Asterisks indicate significant differences between strains (Wilcoxon signed-rank test; p-value 0.05). (B) Boxplot showing the growth of two phytopathogenic bacteria under control conditions (grey) or after exposure to culture filtrates of strain I1237 (pink) and *T. gamsii* T035 (blue). Different letters indicate significant differences according to Tukey’s multiple comparisons test (α = 0.05).

An intriguing pattern emerged in the parasitism assays with *Globisporangium* species: strain T035 was particularly effective against isolate PAT-018, but showed markedly reduced activity against the two other *Globisporangium* strains tested (Fig. 4A).

### Genome sequencing and assembly results

ONT sequencing of *T. gamsii* strain T035 generated 196,604 reads with a N50 read length of 6,669 bp, yielding approximately 965 Mbp of sequence data, which corresponds to around 25X genome coverage (Fig. 5). The median read quality and length were respectively 21.8 and 3,217 bp.

**Figure 5.**
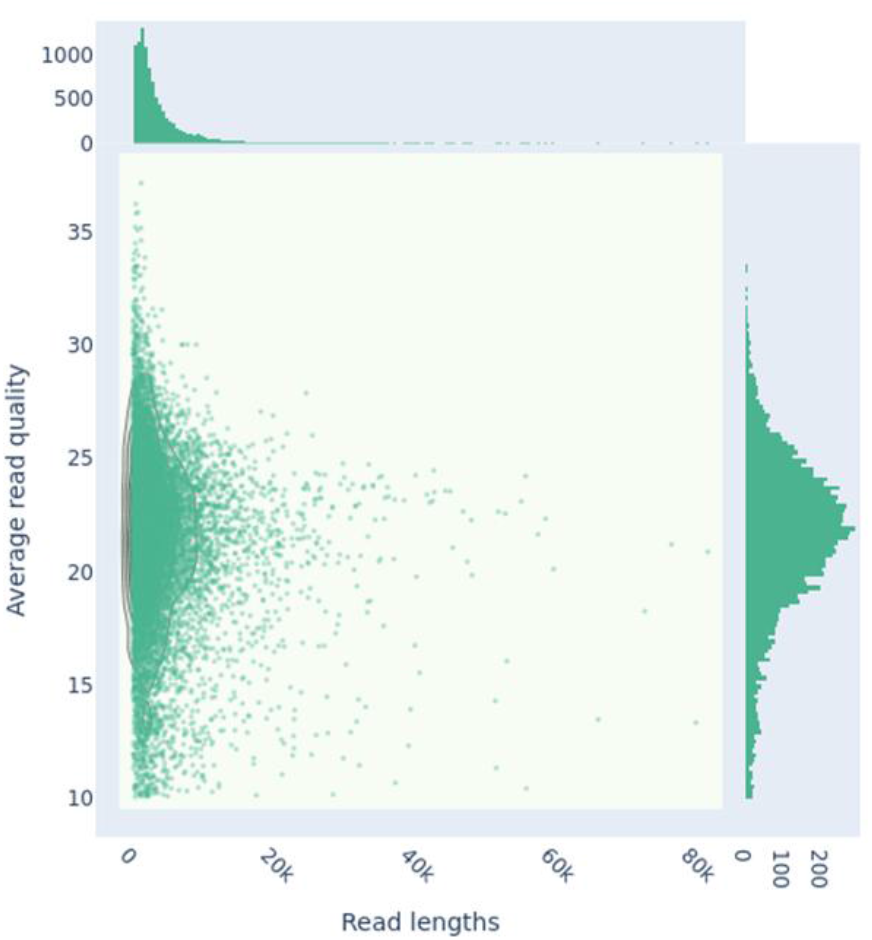
Scatterplot of Nanopore reads for *T. gamsii* strain T035, showing read length versus quality score, generated with NanoPlot (kde plot). A total of 196,604 reads were analyzed, representing approximately 965 Mbp of sequence data. The read length N50 is 6,669 bp, with a median read length of 3,217 bp, while the median read quality is 21.8.

The initial Flye assembly of *T. gamsii* T035 produced 30 contigs (N50 = 3.1 Mbp and N90 = 1.3 Mbp), which were scaffolded into 16 sequences (N50 = 7.18 Mbp and N90 = 4.03 Mbp; 7 nuclear chromosomes, 5 unassigned contigs and 4 mitochondrial sequences), which introduced 3.09 N’s per 100 kbp (Fig. 3, Fig. 6 and Table 1). The total assembly length is of 38.8 Mbp, with a GC content of 48.44% and a largest contig of 7.09 Mbp. Assembly completeness, assessed with BUSCO, indicated 98.8% complete universal genes (complete and single-copy: 98.3%; complete and duplicated: 0.5%; fragmented: 0.4%; missing: 0.8%) against the fungi_odb10 database (Fig. 3) (Tegenfeldt *et al*., 2025). Key genome metrics for T035 are compared with other published *T. gamsii* genomes in Table 1, and synteny analysis between T035, T6085, and A5MH highlights the high fragmentation of T6085 and A5MH relative to T035 (Supplementary Figure S2).

**Figure 6.**
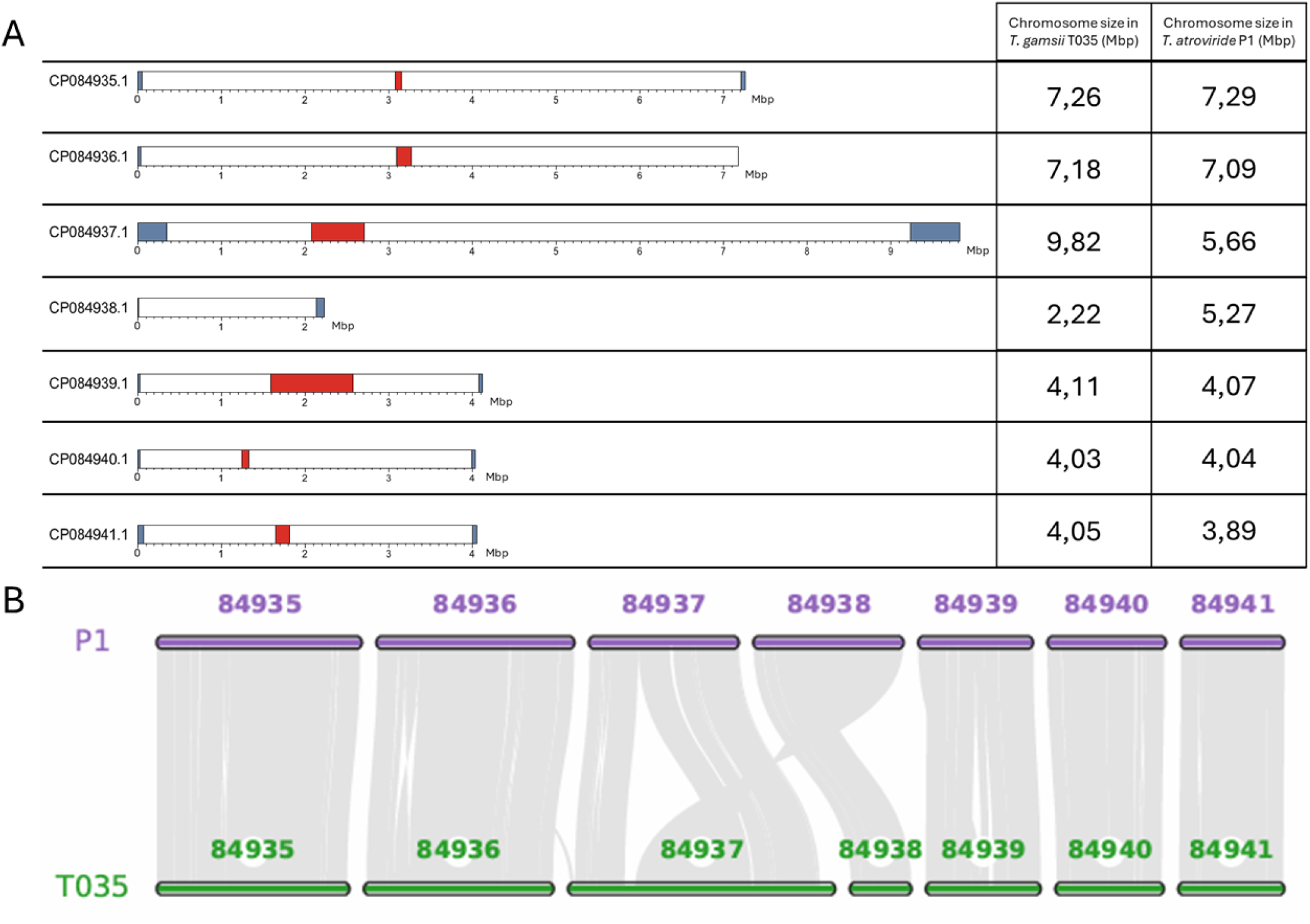
Complete sequences of the seven chromosomes of *T. gamsii* obtained in this study, compared with the reference genome of *T. atroviride* P1. (A) Chromosomal structures of *T. gamsii*. Telomeres are shown in blue and centromeres in red. One telomere was not detected on chromosome CP084936.1, and the centromere was not detected on chromosome CP084938.1. Chromosome sizes for *T. gamsii* T035 and the *T. atroviride* reference genome P1 are shown on the right. (B) Synteny analysis between the genomes of *T. gamsii* T035 and *T. atroviride* P1 shows that the expansion of chromosome CP084937 in *T. gamsii* T035 is due to an interchromosomal translocation from chromosome CP084938.

### Genomic structure

The *T. gamsii* T035 genome is composed of seven large sequences corresponding to chromosomes. These chromosomes are well defined, as most of them are flanked by telomeric regions, except for chromosome CP084936.1 for which one telomeric region was missing (Fig. 6A). Notably, chromosome CP084937.1 is almost twice as long in *T. gamsii* T035 as in *T. atroviride* P1. This size difference is likely attributable to an interchromosomal translocation from chromosome CP084938.1 to chromosome CP084937.1 (Fig. 6B). As a consequence, chromosome CP084938.1 is markedly shorter in *T. gamsii* T035 than in *T. atroviride* P1 (2.22 Mbp versus 5.27 Mbp). This reduction in size may explain why no centromeric region was identified on chromosome CP084938.1 in *T. gamsii* T035 (Fig. 6A).

### Genome structural annotation and annotation quality control

Structural annotation was performed using the deep-learning tool Helixer (Holst *et al*., 2023), which predicted 13,036 protein-coding genes. Compared to the previously published reference genome of *T. gamsii* strain T6085, annotated using MAKER2 (Holt & Yandell, 2011), the annotation of strain T035 identified more than 2,000 additional protein-coding genes (Table 2).

**Table 2.**
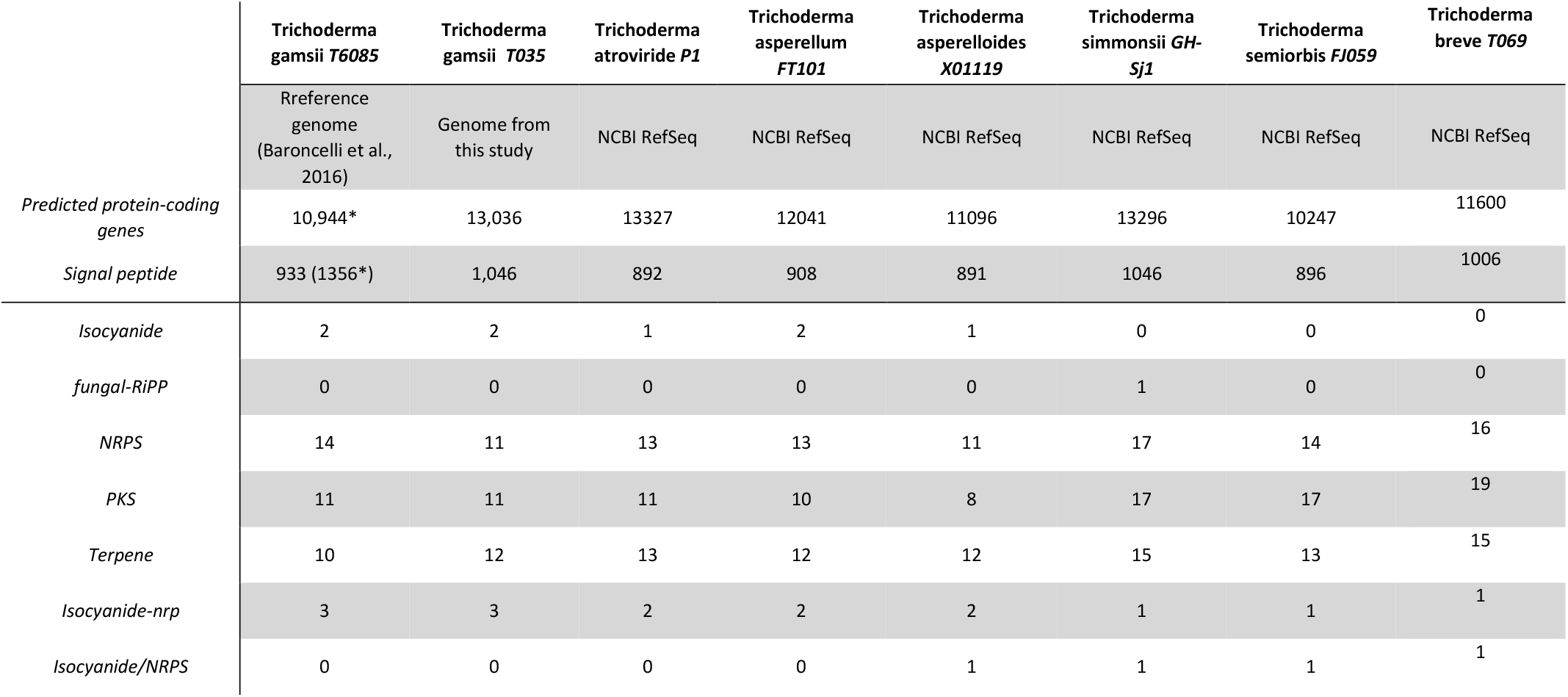
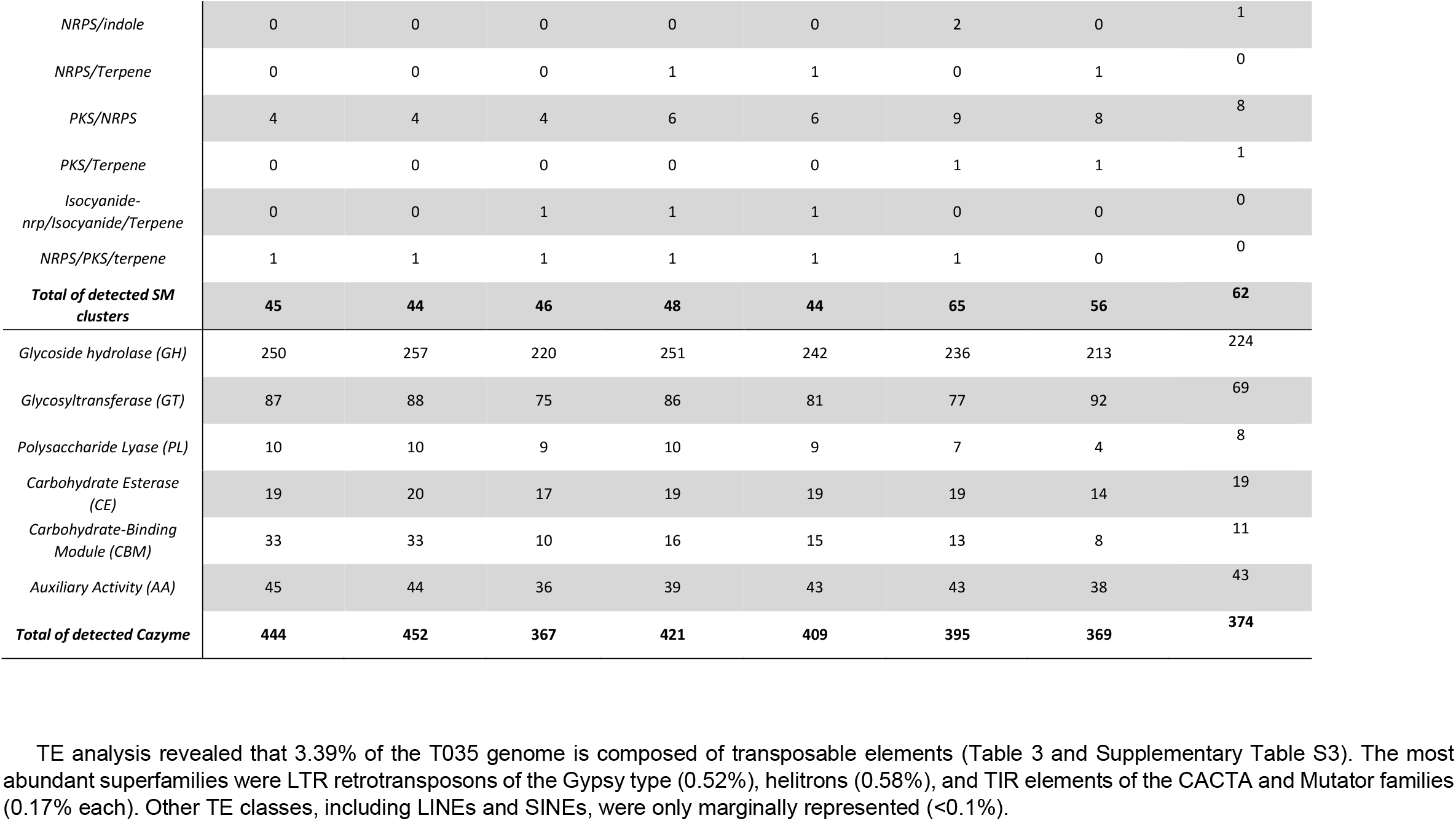
Genome annotation metrics comparing *T. gamsii* (strain T035) from this study with the reference genome *T. gamsii* (strain T6085) from Baroncelli et al. (2016) as well as with the completed genomes available on NCBI. Asterisks indicate values reported in Baroncelli et al. (2016); values without asterisks were generated in this study. Signal peptides were detected using DeepSig, SM clusters were predicted using antiSMASH, and CAZymes were identified using dbCAN3.

Recent studies have reported that deep-learning-based predictors may occasionally generate unsupported gene models (Lapalu *et al*., 2025). Therefore, part of the increase in predicted gene number may reflect differences between annotation pipelines rather than true biological differences between strains.

**Table 3.** Composition of transposable elements in *Trichoderma gamsii* strain T035 and in *Trichoderma atroviride* as determined with EDTA, RIP sequences were identified using The RIPper.

| Class |  | T. gamsii T035 |  |  | T. atroviride P1 |  |  |
| --- | --- | --- | --- | --- | --- | --- | --- |
|  |  | Nb of elements | Nb of RIP | % of sequence | Nb of elements | Nb of RIP | % of sequence |
| LINE | Tad1 | 51 | 0 | 0.06% | 0 | 0 | 0.00% |
| LTR | Gypsy | 181 | 26 | 0.52% | 172 | 49 | 1.06% |
|  | Copia | 0 | 0 | 0.00% | 1 | 0 | 0.00% |
|  | Unknown | 78 | 15 | 0.26% | 9 | 0 | 0.00% |
| SINE | tRNA | 17 | 0 | 0.00% | 19 | 0 | 0.01% |
|  | unknown | 15 | 1 | 0.00% | 0 | 0 | 0.00% |
| TIR | CACTA | 28 | 1 | 0.17% | 25 | 0 | 0.17% |
|  | Mutator | 33 | 0 | 0.17% | 34 | 2 | 0.19% |
|  | PIF_Harbinger | 8 | 0 | 0.03% | 2 | 1 | 0.01% |
|  | Tc1_Mariner | 4 | 0 | 0.03% | 2 | 0 | 0.02% |
|  | hAT | 1 | 0 | 0.00% | 3 | 0 | 0.01% |
| nonTIR | helitron | 23 | 1 | 0.58% | 9 | 0 | 0.24% |
| rDNA | 45S | 13 | 0 | 0.01% | 0 | 0 | 0.00% |
| Repeat_fragment | - | 813 | 46 | 1.56% | 297 | 8 | 0.15% |
| Total |  | 1265 | 90 | 3.39% | 573 | 60 | 1.87% |

### Genome functional annotation

Functional annotation using EggNOG-mapper (Cantalapiedra *et al*., 2021) assigned functional annotations to 9,344 out of 13,036 predicted genes. The mating-type locus of *T. gamsii* T035 was identified using BLAST searches against known MAT proteins. Strong homology was detected with *Trichoderma spinulosum* MAT1-1-1, MAT1-1-2, and MAT1-1-3 protein sequences (best BLAST hits: 429 bits, e-value 1e-121; 549 bits, e-value 1e-157; and 271 bits, e-value 8e-074, respectively). No MAT1-2 genes were detected, indicating that *T. gamsii* T035 carries the MAT1-1 mating type.

Secretome prediction identified 1,046 proteins with signal peptides using DeepSig (Savojardo *et al*., 2018) (Table 2). For comparison, 933 signal peptide-containing proteins were identified using the same tool in the strain T6085, while 1,356 were previously reported using SignalP (Petersen *et al*., 2011; Baroncelli *et al*., 2016). Among the seven *Trichoderma* genomes analyzed in this study, *T. gamsii* T035 exhibited one of the highest number of predicted secreted proteins (Table 2).

SM biosynthetic gene clusters were predicted in both *T. gamsii* strains T035 and T6085, with 44 and 45 clusters identified, respectively (Table 2). The clusters comprised a globally similar distribution of types, including NRPS, PKS, terpene, isocyanide, and mixed NRPS/PKS clusters, reflecting comparable secondary metabolic potential in both strains. Notably, *T. gamsii* harbored fewer SM clusters than some other *Trichoderma* species, *T. simmonsii* in particular exhibited as high as 65 predicted SM clusters (Table 2).

Finally, CAZymes were identified with high confidence based on concordant predictions from HMMER, DIAMOND, and dbCAN_sub. A total of 452 and 444 CAZyme activities were detected in *T. gamsii* strains T035 and T6085, respectively (Table 2). Interestingly, compared with the other analyzed *Trichoderma* species, *T. gamsii* exhibited the highest number of predicted carbohydrate-active enzymes, whereas the other strains ranged from 367 to 421 CAZyme activities. This difference mainly resulted from an overrepresentation of carbohydrate-binding module (CBM) domains, which were two-to three-fold more abundant in *T. gamsii* than in the other tested *Trichoderma* species.

## Discussion

In this study, we generated the most contiguous genome assembly of the biocontrol fungus *T. gamsii* to date, with scaffold N50 and L50 values of 7.18 Mbp and 3, respectively, compared to the previously best available assembly, which had an N50 of 697.4 kbp and an L50 of 18 (Table 2, Fig. 3). The high quality of the assembly and annotation was further supported by a BUSCO analysis (Fig. 3).

Biocontrol assays indicate that *T. gamsii* T035 exhibits higher antagonistic potential than a commercial *Trichoderma* strain, with significant inhibitory activity against several fungal and bacterial pathogens (Fig. 4). The activity is particularly strong against the Basidiomycete *Rhizoctonia solani* and the Ascomycete *Stagonosporopsis* spp., whereas inhibition of wood-decay fungi is minimal. Activity against *Globisporangium* was isolate-dependent, suggesting a degree of host specificity (Fig. 4A).

This specificity could involve host-selective toxins, carbohydrate-active enzymes, and/or secreted effectors, as reported for plant-pathogenic fungi (Hatta *et al*., 2002; Lyu *et al*., 2015; Ayukawa *et al*., 2021; Courtial *et al*., 2022). In *Trichoderma*, host-dependent regulatory pathways have also been described (Sharma *et al*., 2017), supporting the idea that T035’s biocontrol mechanisms may involve specialized attack pathways or differential interactions with host defenses. We therefore speculate that the observed specificity of T035 against *Globisporangium* spp. could involve specialized attack pathways, possibly mediated by host-specific toxins and/or effectors (Ramírez-Valdespino, Casas-Flores et Olmedo-Monfil, 2019), or alternatively reflect differences in host defense mechanisms.

Interestingly, compared to other *Trichoderma* genomes, *T. gamsii* genomes seemed depleted in SM cluster but enriched in secreted proteins and CAZymes, particularly CBM-containing enzymes, which may indicate distinct antagonistic strategies among *Trichoderma* species. CBM-containing enzymes bind to cellulose and chitin and have been reported to be frequently located adjacent to chitinase genes in fungal genomes (Kubicek *et al*., 2011). These CBM domains are linked to the catalytic core of enzymes and enhance their efficiency, especially when acting on recalcitrant substrates or substrates present at low concentrations (Kubicek et Kubicek, 2016). Functional validation of the identified secondary metabolite clusters, CAZymes, and predicted secretome will be required to test these hypotheses.

Additionally, analysis of the global genome structure of *T. gamsii* T035 suggests an interchromosomal translocation from chromosome CP084938.1 to CP084937.1 when compared with *T. atroviride* P1 (Fig. 6B). Confirmation of this rearrangement in other complete *T. gamsii* genomes will help determine when this event could have occurred. Furthermore, the TE content of T035 is relatively low compared to other filamentous fungi, where TE content can go from 0,03% up to more than 80% (e.g., *Fusarium graminearum* and *Blumeria graminis*, respectively) (Ma *et al*., 2010; Kusch *et al*., 2024). However, *Trichoderma* species are generally TE-poor, with major TE classes typically representing 0.48 to 0.57% of the genome (Kubicek *et al*., 2011). In our study, TE content in *T. atroviride* was around 1.9%, while previous estimate for other *T. atroviride* genome was 0.49% (Kubicek *et al*., 2011). By comparison, *T. gamsii* T035 has a higher TE content (3.39%), most of which consists of highly degenerated repeat fragments. Moreover, the proportion of RIP-affected sequences is also higher in *T. gamsii* 93 T035 (2.66%) than in *T. atroviride* P1 (1.77%).

Further studies across multiple *Trichoderma* species and strains will be required to better understand TE dynamics in this genus, as TE content is known to be highly strain-specific, as demonstrated in *Leptosphaeria maculans* (2.7–32.5%) (Grandaubert *et al*., 2014).

Overall, this study provides a high-quality genome assembly of *T. gamsii* T035, including detailed annotation of protein-coding genes, secondary metabolite clusters, carbohydrate-active enzymes, and the mating-type locus (MAT1-1). The T035 genome encodes 13,036 protein-coding genes, compared to 10,944 in the published reference genome of strain T6085. The genome has been deposited in ENA under BioProject PRJEB107169. This genomic resource, combined with functional assays, will support future studies aimed at elucidating the molecular basis of *T. gamsii* biocontrol activity and its potential applications in sustainable agriculture.

## Acknowledgements

We thank the ANAN platform and the entire Nanopore Team, especially Muriel Bahut, Jessica Dittmer, Maryline Cournol, and Sandrine Balzergue, for the material and technical support during the genome sequencing. We acknowledge the GenOuest bioinformatics core facility (https://www.genouest.org) for providing the computing infrastructure.

Preprint version 4 of this article has been peer-reviewed and recommended by Peer Community In Genomics (https://doi.org/10.24072/pci.genomics.100458; Malagnac, 2026).

## Data, scripts, code, and supplementary information availability

Data, supplementary figures and tables, scripts, and code are available online on GitHub and stored with Zenodo: https://doi.org/10.5281/zenodo.18417586 (JustiCol, 2026).

T035 genomic data are available at http://www.ebi.ac.uk/ena/browser/view/PRJEB107169.

## References

Alonge, M. et al. (2022) « Automated assembly scaffolding using RagTag elevates a new tomato system for high-throughput genome editing », Genome Biology, 23(1), p. 258. Disponible sur: 10.1186/s13059-022-02823-7.

Anees, M. et al. (2010) « Characterization of field isolates of Trichoderma antagonistic against Rhizoctonia solani », Fungal Biology, 114(9), p. 691–701. Disponible sur: 10.1016/j.funbio.2010.05.007.

Ayukawa, Y. et al. (2021) « A pair of effectors encoded on a conditionally dispensable chromosome of Fusarium oxysporum suppress host-specific immunity », Communications Biology, 4(1), p. 707. Disponible sur: 10.1038/s42003-021-02245-4.

Baroncelli, R. et al. (2016) « Draft Whole-Genome Sequence of Trichoderma gamsii T6085, a Promising Biocontrol Agent of Fusarium Head Blight on Wheat », Genome Announcements, 4(1), p. e01747–15. Disponible sur: 10.1128/genomeA.01747-15.

Bedine Boat, M.A. et al. (2020) « Screening, identification and evaluation of Trichoderma spp. for biocontrol potential of common bean damping-off pathogens », Biocontrol Science and Technology, 30(3), p. 228–242. Disponible sur: 10.1080/09583157.2019.1700909.

Blin, K. et al. (2025) « antiSMASH 8.0: extended gene cluster detection capabilities and analyses of chemistry, enzymology, and regulation », Nucleic Acids Research, 53(W1), p. W32–W38. Disponible sur: 10.1093/nar/gkaf334.

Bunbury-Blanchette, A.L. et Walker, A.K. (2019) « Trichoderma species show biocontrol potential in dual culture and greenhouse bioassays against Fusarium basal rot of onion », Biological Control, 130, p. 127–135. Disponible sur: 10.1016/j.biocontrol.2018.11.007.

Cantalapiedra, C.P. et al. (2021) « eggNOG-mapper v2: Functional Annotation, Orthology Assignments, and Domain Prediction at the Metagenomic Scale », Molecular Biology and Evolution. Édité par K. Tamura, 38(12), p. 5825–5829. Disponible sur: 10.1093/molbev/msab293.

Chateau, C. et al. (2024) « In vitro and in vivo biocontrol studies against Botrytis cinerea, the causal agent of grey mould in Hydrangea macrophylla », Biocontrol Science and Technology, 34(3), p. 269–295. Disponible sur: 10.1080/09583157.2024.2341302.

Chen, J.-L. et al. (2016) « Endophytic Trichoderma gamsii YIM PH30019: a promising biocontrol agent with hyperosmolar, mycoparasitism, and antagonistic activities of induced volatile organic compounds on root-rot pathogenic fungi of Panax notoginseng », Journal of Ginseng Research, 40(4), p. 315–324. Disponible sur: 10.1016/j.jgr.2015.09.006.

Courtial, J. et al. (2022) « Characterization of NRPS and PKS genes involved in the biosynthesis of SMs in Alternaria dauci including the phytotoxic polyketide aldaulactone », Scientific Reports, 12(1), p. 8155. Disponible sur: 10.1038/s41598-022-11896-0.

De Coster, W. et al. (2018) « NanoPack: visualizing and processing long-read sequencing data », Bioinformatics. Édité par B. Berger, 34(15), p. 2666–2669. Disponible sur: 10.1093/bioinformatics/bty149.

Dennis, C. et Webster, J. (1971) « Antagonistic properties of species-groups of Trichoderma », Transactions of the British Mycological Society, 57(1), p. 25–IN3. Disponible sur: 10.1016/S0007-1536(71)80077-3.

Galletti, S., Paris, R. et Cianchetta, S. (2020) « Selected isolates of Trichoderma gamsii induce different pathways of systemic resistance in maize upon Fusarium verticillioides challenge », Microbiological Research, 233, p. 126406. Disponible sur: 10.1016/j.micres.2019.126406.

Grandaubert, J. et al. (2014) « Transposable element-assisted evolution and adaptation to host plant within the Leptosphaeria maculans-Leptosphaeria biglobosa species complex of fungal pathogens », BMC Genomics, 15(1), p. 891. Disponible sur: 10.1186/1471-2164-15-891.

Hatta, R. et al. (2002) « A Conditionally Dispensable Chromosome Controls Host-Specific Pathogenicity in the Fungal Plant Pathogen Alternaria alternata », Genetics, 161(1), p. 59–70. Disponible sur: 10.1093/genetics/161.1.59.

Hofstatter, P.G. et al. (2022) « Repeat-based holocentromeres influence genome architecture and karyotype evolution », Cell, 185(17), p. 3153-3168.e18. Disponible sur: 10.1016/j.cell.2022.06.045.

Holst, F. et al. (2023) « Helixer– de novo Prediction of Primary Eukaryotic Gene Models Combining Deep Learning and a Hidden Markov Model ». Bioinformatics. Disponible sur: 10.1101/2023.02.06.527280.

Joubert, A. et al. (2010) « Laser nephelometry applied in an automated microplate system to study filamentous fungus growth », BioTechniques, 48(5), p. 399–404. Disponible sur: 10.2144/000113399.

JustiCol (2026) « JustiCol/Genome-assembly-T_gamsii_T035: T_gamsii_T035_genome_R1.2 ». Zenodo. Disponible sur: 10.5281/ZENODO.18417586.

Katz, L. et al. (2019) « Mashtree: a rapid comparison of whole genome sequence files », Journal of Open Source Software, 4(44), p. 1762. Disponible sur: 10.21105/joss.01762.

Kolmogorov, M. et al. (2019) « Assembly of long, error-prone reads using repeat graphs », Nature Biotechnology, 37(5), p. 540–546. Disponible sur: 10.1038/s41587-019-0072-8.

Koren, S. et al. (2017) « Canu: scalable and accurate long-read assembly via adaptive k -mer weighting and repeat separation », Genome Research, 27(5), p. 722–736. Disponible sur: 10.1101/gr.215087.116.

Kubicek, C.P. et al. (2011) « Comparative genome sequence analysis underscores mycoparasitism as the ancestral life style of Trichoderma », Genome Biology, 12(4), p. R40. Disponible sur: 10.1186/gb-2011-12-4-r40.

Kubicek, C.P. et Kubicek, E.M. (2016) « Enzymatic deconstruction of plant biomass by fungal enzymes », Current Opinion in Chemical Biology, 35, p. 51–57. Disponible sur: 10.1016/j.cbpa.2016.08.028.

Kumar, S. et al. (2024) « MEGA12: Molecular Evolutionary Genetic Analysis Version 12 for Adaptive and Green Computing », Molecular Biology and Evolution. Édité par F.U. Battistuzzi, 41(12), p. msae263. Disponible sur: 10.1093/molbev/msae263.

Kusch, S. et al. (2024) « Long-term and rapid evolution in powdery mildew fungi », Molecular Ecology, 33(10), p. e16909. Disponible sur: 10.1111/mec.16909.

Lapalu, N. et al. (2025) « Improved Gene Annotation of the Fungal Wheat Pathogen Zymoseptoria tritici Based on Combined Iso-Seq and RNA-Seq Evidence », Molecular Plant-Microbe Interactions®, p. MPMI-07-25-0077-TA. Disponible sur: 10.1094/MPMI-07-25-0077-TA.

Lee, J. et al. (2012) « The antagonistic properties of Trichoderma spp. inhabiting woods for potential biological control of wood-damaging fungi », Holzforschung, 66(7), p. 883–887. Disponible sur: 10.1515/hf.2011.187.

Li, W.-C. et al. (2021) « Complete Genome Sequences and Genome-Wide Characterization of Trichoderma Biocontrol Agents Provide New Insights into their Evolution and Variation in Genome Organization, Sexual Development, and Fungal-Plant Interactions », Microbiology Spectrum. Éditépar C.A. Cuomo, 9(3), p. e00663–21. Disponible sur: 10.1128/Spectrum.00663-21.

Lu, H., Giordano, F. et Ning, Z. (2016) « Oxford Nanopore MinION Sequencing and Genome Assembly », Genomics, Proteomics & Bioinformatics, 14(5), p. 265–279. Disponible sur: 10.1016/j.gpb.2016.05.004.

Lyu, X. et al. (2015) « Comparative genomic and transcriptional analyses of the carbohydrate-active enzymes and secretomes of phytopathogenic fungi reveal their significant roles during infection and development », Scientific Reports, 5(1), p. 15565. Disponible sur: 10.1038/srep15565.

Ma, L.-J. et al. (2010) « Comparative genomics reveals mobile pathogenicity chromosomes in Fusarium », Nature, 464(7287), p. 367–373. Disponible sur: 10.1038/nature08850.

Malagnac, F. (éd.) (2026) « Recommendation of: Genome of Trichoderma gamsii strain T035 a promising beneficial fungus in agriculture. Round#3 », Peer Community in Genomics, p. genomics.100458. Disponible sur: 10.24072/pci.genomics.100458.

Marçais, G. et Kingsford, C. (2011) « A fast, lock-free approach for efficient parallel counting of occurrences of k -mers », Bioinformatics, 27(6), p. 764–770. Disponible sur: 10.1093/bioinformatics/btr011.

Marik, T. et al. (2025) « Novel peptaibiotics identified from Trichoderma clade Viride », Natural Products and Bioprospecting, 15(1), p. 48. Disponible sur: 10.1007/s13659-025-00524-9.

Migheli, Q. et al. (2009) « Soils of a Mediterranean hot spot of biodiversity and endemism (Sardinia, Tyrrhenian Islands) are inhabited by pan-European, invasive species of Hypocrea/Trichoderma », Environmental Microbiology, 11(1), p. 35–46. Disponible sur: 10.1111/j.1462-2920.2008.01736.x.

Mikheenko, A. et al. (2023) « WebQUAST: online evaluation of genome assemblies », Nucleic Acids Research, 51(W1), p. W601–W606. Disponible sur: 10.1093/nar/gkad406.

Möller, E.M. et al. (1992) « A simple and efficient protocol for isolation of high molecular weight DNA from filamentous fungi, fruit bodies, and infected plant tissues », Nucleic Acids Research, 20(22), p. 6115–6116. Disponible sur: 10.1093/nar/20.22.6115.

Pascouau, C. et al. (2023) « Characterization and pathogenicity of Fusarium spp. isolates causing root and collar rot on carrot », Canadian Journal of Plant Pathology, 45(1), p. 76–91. Disponible sur: 10.1080/07060661.2022.2103737.

Petersen, T.N. et al. (2011) « SignalP 4.0: discriminating signal peptides from transmembrane regions », Nature Methods, 8(10), p. 785–786. Disponible sur: 10.1038/nmeth.1701.

Ramírez-Valdespino, C.A., Casas-Flores, S. et Olmedo-Monfil, V. (2019) « Trichoderma as a Model to Study Effector-Like Molecules », Frontiers in Microbiology, 10, p. 1030. Disponible sur: 10.3389/fmicb.2019.01030.

Ranallo-Benavidez, T.R., Jaron, K.S. et Schatz, M.C. (2020) « GenomeScope 2.0 and Smudgeplot for reference-free profiling of polyploid genomes », Nature Communications, 11(1), p. 1432. Disponible sur: 10.1038/s41467-020-14998-3.

Rinu, K., Sati, P. et Pandey, A. (2014) « Trichoderma gamsii (NFCCI 2177): A newly isolated endophytic, psychrotolerant, plant growth promoting, and antagonistic fungal strain », Journal of Basic Microbiology, 54(5), p. 408–417. Disponible sur: 10.1002/jobm.201200579.

Rosolen, R.R. et al. (2022) « Whole-Genome Sequencing and Comparative Genomic Analysis of Potential Biotechnological Strains of Trichoderma harzianum, Trichoderma atroviride, and Trichoderma reesei ». Genomics. Disponible sur: 10.1101/2022.02.11.479986.

Savojardo, C. et al. (2018) « DeepSig: deep learning improves signal peptide detection in proteins », Bioinformatics. Édité par A. Valencia, 34(10), p. 1690–1696. Disponible sur: 10.1093/bioinformatics/btx818.

Schalamun, M. et Schmoll, M. (2022) « Trichoderma – genomes and genomics as treasure troves for research towards biology, biotechnology and agriculture », Frontiers in Fungal Biology, 3, p. 1002161. Disponible sur: 10.3389/ffunb.2022.1002161.

Sharma, V. et al. (2017) « Elucidation of biocontrol mechanisms of Trichoderma harzianum against different plant fungal pathogens: Universal yet host specific response », International Journal of Biological Macromolecules, 95, p. 72–79. Disponible sur: 10.1016/j.ijbiomac.2016.11.042.

Su, W. et al. (2021) « A Tutorial of EDTA: Extensive De Novo TE Annotator », in J. Cho (éd.) Plant Transposable Elements. New York, NY: Springer US (Methods in Molecular Biology), p. 55–67. Disponible sur: 10.1007/978-1-0716-1134-0_4.

Tang, H. et al. (2024) « JCVI: A versatile toolkit for comparative genomics analysis », iMeta, 3(4), p. e211. Disponible sur: 10.1002/imt2.211.

Tegenfeldt, F. et al. (2025) « OrthoDB and BUSCO update: annotation of orthologs with wider sampling of genomes », Nucleic Acids Research, 53(D1), p. D516–D522. Disponible sur: 10.1093/nar/gkae987.

Valan Arasu, M. et al. (2023) « Biocontrol of Trichoderma gamsii induces soil suppressive and growth-promoting impacts and rot disease-protecting activities », Journal of Basic Microbiology, 63(7), p. 801–813. Disponible sur: 10.1002/jobm.202300016.

Vaser, R. et al. (2017) « Fast and accurate de novo genome assembly from long uncorrected reads ». Disponible sur: 10.1101/gr.214270.116.

Warringer, J. et Blomberg, A. (2003) « Automated screening in environmental arrays allows analysis of quantitative phenotypic profiles in Saccharomyces cerevisiae », Yeast, 20(1), p. 53–67. Disponible sur: 10.1002/yea.931.

Zheng, J. et al. (2023) « dbCAN3: automated carbohydrate-active enzyme and substrate annotation », Nucleic Acids Research, 51(W1), p. W115–W121. Disponible sur: 10.1093/nar/gkad328.

